# Hypoxia modulates cellular endocytic pathways and organelles with enhanced cell migration and 3D cell invasion

**DOI:** 10.1101/2022.02.16.480665

**Authors:** Hema Naveena A, Dhiraj Bhatia

**Affiliations:** Biological Engineering Discipline, Indian Institute of Technology Gandhinagar, Palaj Gandhinagar 382355, Gujarat, India

**Keywords:** Cancer hypoxia, cellular endocytosis, cell invasion and migration, cell proliferation

## Abstract

Hypoxia, a decrease in cellular or tissue level oxygen content, is characteristic of most tumours and shown to drive cancer progression by altering multiple subcellular processes. We hypothesized that the cancer cells in a hypoxic environment might have slower proliferation rates and increased invasion and migration rate with altered endocytosis when compared to the cancer cells in the periphery of the tumour mass that experiences normoxic condition. Using chemically induced hypoxia, a short hypoxic exposure increased the uptake of clathrin independent endocytic marker Galectin-3, but a prolonged hypoxic exposure decreased clathrin-independent endocytic uptake, while clathrin mediated endocytosis remained unaffected. Subcellular organelles such as mitochondria showed enhanced intensity to withstand the hypoxic stress, while other organelles such as ER were significantly decreased. The proliferation rates decreased, and the migration and invasion rate increased in cancer cells in hypoxic condition compared to normoxic cancer cells.These data suggest that hypoxia modulates cellular endocytic pathways with decreased proliferation and enhanced cell migration and invasion.

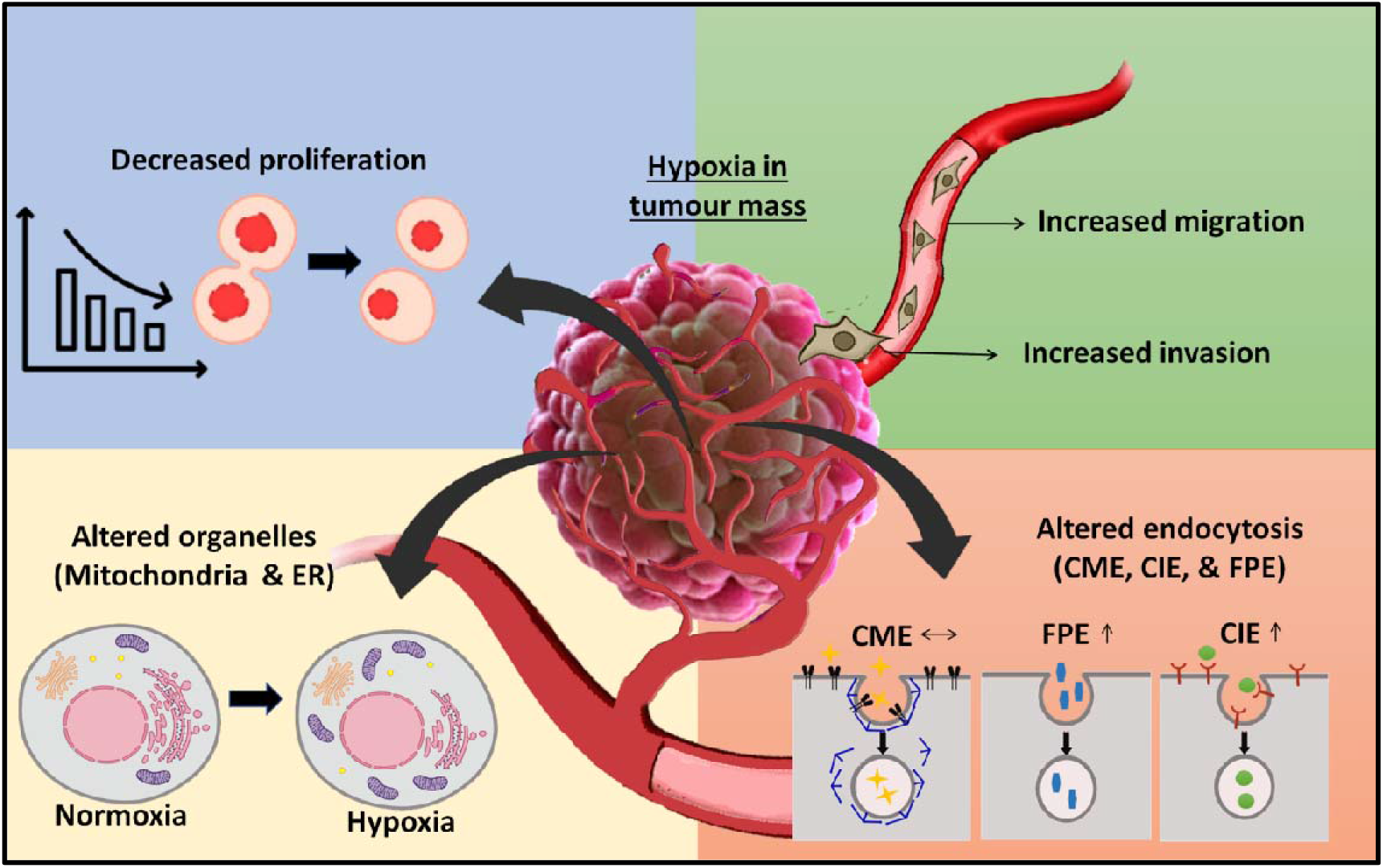

## Introduction

Cancer is a very complex condition of cells and tissues, and it is the result of several successive and collective cellular changes (1). During the tumour development and progression, the cancer cells and the other cells in the tumour microenvironment have limited access to nutrients and oxygen, especially in the core of the tumour tissue, which is at a distance from the tumour induced vasculature (2, 3). Such a region with a low oxygen level is termed a hypoxic region. The distribution of hypoxic levels in tumour tissue is highly heterogeneous, and it ranges from mild hypoxic to severe anoxic conditions(4). Hypoxia induces several molecular events that favor cancer progression by forming aggressive clones from heterogeneous cancer cell populations, leading to lethal phenotypes(5). This condition in tumour microenvironment leads to poor prognosis and poor outcomes of anti-cancer therapies(6).

Cancer cells’ response to the hypoxic condition is ascribed to hypoxic inducible factor (HIF). HIF has two subunits; O_2_ dependent α subunit (HIFα), which dimerize with β subunit (HIFβ) that are constitutively expressed(7). In the presence of O_2_, the two proline residues in theα subunits are hydroxylated by prolyl hydroxylase enzyme that promotes binding to von Hippel-Lindau protein, which accelerates HIFα ubiquitination and degradation(8, 9). Both subunits dimerize to form an active transcription factor in hypoxic condition and switch the cell metabolism to glycolytic mode with increased glucose consumption and increased pyruvate, lactate, and H+ production. These transcriptional responses driven by HIF upon initiation of hypoxia can promote the adaptation and selection of cancer cells in comparison to surrounding normoxic condition, thus favoring cancer progression(5).Hypoxia was stimulated in-vitro by incubation with chemical agents such as cobalt chloride, a well-established chemical hypoxic mimicking agent (10)

Hypoxia,a deviation from the homeostatic condition, is generally unfavorable for cells and tissues, but the cells in the tumour microenvironment adapt to cope with the depleted oxygen levels. Plasma membrane would play an essential role in adaptation to hypoxic stress because it is a source of interaction between the cell and their environment through various transmembrane proteins such as transporter, channels, and receptors(11). It adapts to stress by dynamic remodeling of cell surface proteins mediated by multiple endocytic pathways (12–14). Though different molecular aspects during hypoxic stress are explored, how different endocytic pathways and the resultant cargo uptake are altered during hypoxic conditions remains unexplored.

We attempt to partly answer some of the fundamental questions in cell biology about hypoxia: a) How does hypoxia affect the cellular organelles’organization and endocytic pathways? b) Does hypoxia affects the proliferation and migration rate of cells? c) Is hypoxia’s effect in cellular organelles correlated with proliferation and migration? Our results open new avenues to explore how different endocytic pathways and organelle organization and dynamics are affected in hypoxia. Are there any concerted mechanisms that might respond to hypoxia and lead to changes in cellular physiology? A better understanding of these factors might help further understand the collective response of cells and tissues to conditions like hypoxia and might help future designs of targeted therapeutics.

## Materials and methods

### Materials

Dulbecco’s modified Eagle’s medium (DMEM), fetal bovine serum (FBS), penicillin−streptomycin, trypsin−EDTA (0.25%), phosphate buffer saline (PBS) were purchased from Gibco. Transferrin-A561, phalloidin A488, FITC-dextran 10kDa, MTT (3-(4,5-Dimethylthiazol-2-yl)-2,5-diphenyltetrazolium Bromide), dihydrorhodamine, DAPI were purchased from Sigma. Cell culture dishes for adherent cells (treated surface), Triton X-100 were purchased from Himedia. Petridishes, culture plates, and live-cell Petri dishes were obtained from Tarsons. Paraformaldehyde (PFA) was purchased from Merck. DMSO was ordered from SRL. CoCl_2_ was purchased from Loba Chem. Collagen Type I was purchased from Corning. Mitotracker deep red FM, ERtracker red, and Hoechst were purchased from Thermo Fisher.Fluorophore-labeled Gal3 were provided as kind gifts from the Johannes team at Institut Curie, Paris.

### Cell culture

Human triple-negative cancer line MDA-MB-231 was cultured in Dulbecco modified eagle medium (DMEM) supplemented with 10% FBS and 1% antimycotic-antibiotic (penicillin-streptomycin) solution. The cells were incubated at 37°C in the humidified chamber containing 5% CO_2_. Generally, the cells were passaged once 85-95% confluency was reached and media was changed as required.

### Induction of cancer hypoxic condition and measurement of reactive oxygen species

Hypoxia was induced chemically with cobalt chloride (CoCl_2_). The aqueous media remains oxygenated while CoCl_2_ induces signaling events associated with hypoxia(15). 10mM stock solution of CoCl_2_ in autoclaved water was freshly prepared. Then subsequent working concentrations (25,50,75,100,200, and 400μM) were prepared in DMEM complete media. The induction of hypoxia was confirmed by detecting reactive oxygen species (ROS) with Dihydrorhodamine 123 (DHR). DHR is an uncharged, non-fluorescent dye that passively diffuses the plasma membrane and acts as a ROS indicator. Upon detection of ROS, it is oxidized to cationic rhodamine 123 and localizes to mitochondria, and exhibits green fluorescence(16). The stock DHR solution (5mM) was prepared in DMSO, and the working DHR concentration (2.5μM) was prepared in DMEM serum-free media.

MDA-MB-231 cells were seeded over the coverslips and incubated at 37°C for 24 h. Hypoxia was induced by adding 150μM CoCl_2_ and incubated at 37°C for 24 h. Control sets were without exposure to CoCl_2_. After 24 h, the media was removed, the cells were washed with 1x PBS, and 2.5 μM DHR in DMEM serum-free media was added and incubated at 37°C for 30 min. The cells were washed with 1X PBS thrice and fixed with 4% paraformaldehyde (PFA) at 37°C for 20 mins. The coverslips were mounted on the slides with mowiol and DAPI, the latter to stain the nucleus. They were imaged using confocal microscopy with a 63X oil immersion objective. The ROS generation, which confirms the induction of hypoxia, was visualized by exciting at 488nm and collecting the fluorescence at the emission range of 498-520nm. The nucleus was visualized by exciting at 405nm and collecting the emission in 415-450nm.

### Optimization of hypoxic condition for MDA-Mb-231 cell lines

The study’s principal aim was to induce hypoxia with less cellular death to understand the cellular effects. Hence optimization of hypoxic conditions induced by CoCl_2_ was required. Optimization was carried out by inducing hypoxia with various concentrations of CoCl_2_. The cells were seeded over the coverslips and incubated at 37°C for 24 h. Hypoxia was induced by adding CoCl_2_ in a series of concentrations of 25, 50, 75, 100,and 200μM and incubated at 37°C for 24 h. Control sets were without exposure to CoCl_2_. For one set after 24 h, the media was removed, and the cells were washed with 1X PBS, and 2.5 μM DHR in DMEM serum-free media was added and incubated at 37°C for 30 mins. The cells were washed with 1X PBS thrice and fixed with 4% PFA at 37°C for 20 mins. The coverslips were mounted on the slides with mowiol and DAPI (to stain the nucleus). For another set, after 24 h, the cells were washed with 1X PBS and fixed with 4% PFA at 37°C for 20 mins. The cells were permeabilized with 0.1% Triton X-100 for 15 mins. Then the cells were treated with 0.1% triton x-100 and 0.1% phalloidin to stain the F-actin filaments to visualize the cell area.The coverslips were mounted on the slides with mowiol and DAPI. They were imaged using confocal microscopy with a 63x oil immersion objective. The ROS generation, which confirms the induction of hypoxia, was visualized by exciting at 488nm and collecting the fluorescence at the emission range of 498-520nm (Emission in green region). The phalloidin staining was visualized by exciting at 488nm and collecting the fluorescence at the emission range of 498 – 520 nm. The nucleus was visualized by exciting at 405nm and collecting the emission in 415 – 450 nm.

### Effect of hypoxia induced chemically by CoCl2 on cell viability in MBA-MB-231 cells (MTT assay)

MTT assay was performed to measure the cell viability(17). MDA-MB-231 cells were seeded in 96 well plates at the seeding density of 10,000 cells per well. The culture plates were incubated at 37°C for 24 h. The media was removed, and the cells were exposed to different concentrations of CoCl_2_ (25, 50, 75, 100, 200, and 400μM). Then it was incubated at 37°C for 24 h. The same was repeated for control without exposure of CoCl_2_. After incubation, 0.5mg/ml of MTT (3-(4,5-Dimethylthiazol-2-yl)-2,5-diphenyltetrazolium Bromide)solution was added to each well and incubated at 37°C for 4 h. The solution was removed and replaced with 100μl of dimethyl sulfoxide (DMSO) in each well and incubated in the dark for 15 mins to dissolve the formazan crystal. The multiwell microplate reader was used to measure absorbance at 570 nm. The cell viability percentage was calculated using equation 1 (18).

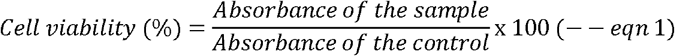

### Effect of hypoxia induced chemically by CoCl_2_ on various endocytic pathways

Endocytosis pathways can be classified as clathrin-mediated pathway (CME) where endocytosis is accomplished by the formation of clathrin-mediated pits and clathrin-independent pathways (CIE) where there is no requirement of clathrin machinery during membrane bending and scission(19). Transferrin was used as an endocytic marker to mark the CME pathway(20) and Galectin-3 (Gal3) to mark the CIE pathway(21). MDA-MB-231 cells were seeded and incubated at 37°C for 24 h, following which hypoxia was induced 100 μM CoCl_2_ for 24 h. The cells were incubated with fluorescently labeled transferrin-A561 and Gal3-A647 (5 μg/mL each) for 30 mins at 37 °C. Then the cells were washed with 1x PBS thrice and fixed with 4% PFA. The fixed cells were mounted on the slides with mowiol and DAPI. In order to study the fluid phase endocytosis (FPE), FITC-dextran 10kDa was used(22). Post induction of hypoxia for 24 h, cells were incubated with FITC-dextran 10kDa (25μg/ml) for 30 mins at 37°C, washed, and fixed. The fixed cells were mounted on the slides with mowiol and DAPI. Confocal imaging was carried out with a 63x oil immersion objective. The transferrin uptake was visualized by exciting at 561nm and collecting the fluorescence at the emission range of 571-620nm. Galectin-3 uptake was visualized by exciting at 633nm and collecting the fluorescence at the emission range of 645-700nm. The nucleus was visualized by exciting at 405nm and collecting the emission in 415-450nm.

### Effect of hypoxia in cellular organelles

Organelles play critical roles in cellular physiology and are altered drastically upon different kinds of stress. Cellular organelles were analyzed at hypoxic stress condition. Different cellular organelles were stained and visualized using published protocols and reagents(23, 24). MDA-MB-231 cells were seeded in live-cell Petri dishes for live-cell imaging organelles. Hypoxia was induced by 100μM of CoCl_2_ for 30 mins on the following day. **(i)** Mitotracker deep red FM was specifically used to stain the mitochondria. A final concentration of 50nM of mitotracker deep red was prepared in DMEM serum-free media and added to Petri dishes containing the cells. They were incubated at 37°C for 30 mins. Further, the cells were washed with 1X PBS and imaged at live-cell condition. **(ii)** Endoplasmic reticulum was stained with ER tracker. A final concentration of 50nM was prepared in DMEM serum-free media, and the cells were incubated at 37°C for 30 mins. The cells were washed with 1X PBS and imaged at live condition. Mitochondria were visualized by exciting at 633nm and collecting the emission range of 645-700nm. Similarly, ER was visualized by exciting at 561nm and collecting the emission signals in the range of 571-620nm.

### Assessment of collective cell migration in normoxic and hypoxic condition with scratch assay

MDA-MB-231 cells were grown to 90% confluency in 35mm petri dishes. A fine scratch was made with a 10μl pipette tip through the center of the MDA-MB-231 monolayer. The cellular debris was removed, and the cell monolayer was washed with 1ml of growth media to smoothen the edge of the scratch(25). The addition of 100μM CoCl_2_ induced hypoxia,and normoxia was maintained without exposure to CoCl_2_. The width of the scratch was measured immediately (0 h) and after an incubation period of 12 h and 24 h under a phase-contrast microscope to determine the rate of cell migration in normoxic and hypoxic conditions. The wound area was recorded immediately (0 h) and after an incubation period of 12 h and 24 h under a phase-contrast microscope to determine the extent of wound closure. The rate of cell migration (R_M_) and the percentage wound closure were calculated according to the following equations 2 and 3 (26).

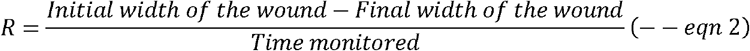

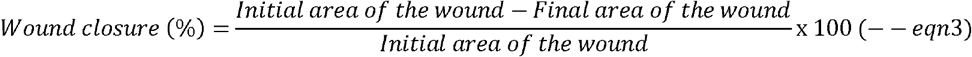

### Effect of hypoxia induced chemically by CoCl_2_ on cell migration 3D spheroid model

Three-dimensional spheroid model using MDA-MB-231 cells is the most effective method to study invasion as they closely mimic the tumour microenvironment. MDA-MB-231 cells were used to form the spheroids by the hanging drop method(27). The cell suspension was placed as a drop underneath the petri plate lids. The cells accumulate at the tip of the drop, aggregate, and form spheroids. Uniform cell suspension of MDA-MB-231 cells in DMEM complete media was taken in a concentration of 5,000 cells/mL. From the homogenous suspension, an aliquot of 20 μl was placed as a drop in the inner surface of the petri plate lid.The petri plate was filled with 1x PBS (~25ml) to maintain a humid environment and prevent droplets from evaporating. The lid consisting of the droplets was gently inverted over the Petri plate and was incubated at 37 °C for 24 h to facilitate 3D spheroid formation. After 24 h, the spheroid formation was visualized using an optical microscope with 10X objectives. After the spheroids were formed, the spheroids were dispersed gently onto the coverslips containing 50μl of collagen-DMEM complete media matrix (in 3:1 vol/vol ratio) in well plates for control sets. The invasion of spheroids in hypoxic condition induced by 100μM and 200μM CoCl_2_ was checked by dispersing the spheroids gently onto the coverslips containing 50μl of collagen-DMEM complete media matrix saturated with 100μM and 200μM CoCl_2_ (in 3:1 v/v ratio) in culture plates. The spheroids placed in the matrix were incubated at 37°C for 1 h. The DMEM complete media was supplemented, and hypoxia was induced by adding 100μM and 200μM CoCl_2_ to DMEM complete media. It was incubated at 37°C for 24 h.

Five spheroids were taken for each control and hypoxic condition.After 24 h, the spheroids were fixed with 4% PFA at 37°C for 20 mins. The spheroids were permeabilized with 0.1% Triton X-100 for 30 mins and stained with 0.1% phalloidin A488 for 1 h to visualize the actin filaments. Followed by staining, it was washed with 1X PBS thrice and mounted on the slides along with the nuclear stain DAPI.

Further, the spheroids were visualized using a confocal microscope with a 10X objective. The excitation wavelength of 405nm and 488nm was used, and the emission was monitored in blue (DAPI, nucleus) and green (Phalloidin and DHR, actin and ROS, respectively) regions. The invasion distance of the tumor cells from the surface of the 3D tumor models was calculated with the ImageJ software. The invasion index of the cells from the spheroid was observed with respect to control, and the invasion index was calculated using the following equation 4 (18).

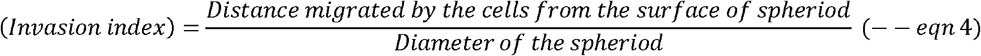

### Effect of hypoxia in cell proliferation

The effect of hypoxia in cell proliferation was studied using MTT and manual cell counting(28). For MTT, approximately 10,000 cells were seeded, and the cells were allowed to attach to the surface.Hypoxia was induced by the addition of 100μM of CoCl_2_ and was incubated at 37°C for 48 h. The control set was maintained without exposure to CoCl_2_. To determine the seeding density with MTT, the cells were maintained with serum-free media to arrest proliferation. The cells were incubated at 37°C for 48 h. After incubation,0.5mg/ml of MTT solution was added to each well and incubated at 37°C for 4 h.The solution was removed and replaced with dimethyl sulfoxide (DMSO) in each well and incubated in the dark for 15 mins to dissolve the formazan crystal. The multiwell microplate reader was used to measure absorbance at 570 nm. For manual counting approximately, 1.5 x 10^5^ cells were seeded, and the cells were allowed to attach to the surface. Hypoxia was induced by the addition of 100μM of CoCl_2_ and was incubated at 37°C for 48 h. The control set was maintained without exposure to CoCl_2_. Then the cells were trypsinized in both conditions and manually counted using a hemocytometer.

### Imaging, image processing, and statistical analysis

Confocal images were captured using Leica confocal microscopy. The live-cell imaging was carried out using the live-cell chamber, which maintains humidity (90%), temperature (37°C), and CO_2_ (5%). The images were processed using Image J 2.1.0. The difference between the control and test groups’ mean values was analyzed by the student’s t-test or one-way ANOVA using GraphPad Prism 9.0.

## Results and discussion

### Induction of hypoxia

Hypoxia was induced chemically with CoCl_2_, and the induction of hypoxia was confirmed by monitoring the ROS species by DHR. **Figure 1A** shows ROS detection by DHR,indicating successful hypoxia induction. The control cells showed a minimum fluorescent signal. The hypoxic cells showed a significant increase in the fluorescent signal,indicating higher ROS generation(29). Thus, it was confirmed that CoCl_2_ induced ROS, which indicates the generation of hypoxia and detection of ROS with DHR, was used for confirming hypoxiafor further studies.

**Figure 1.**
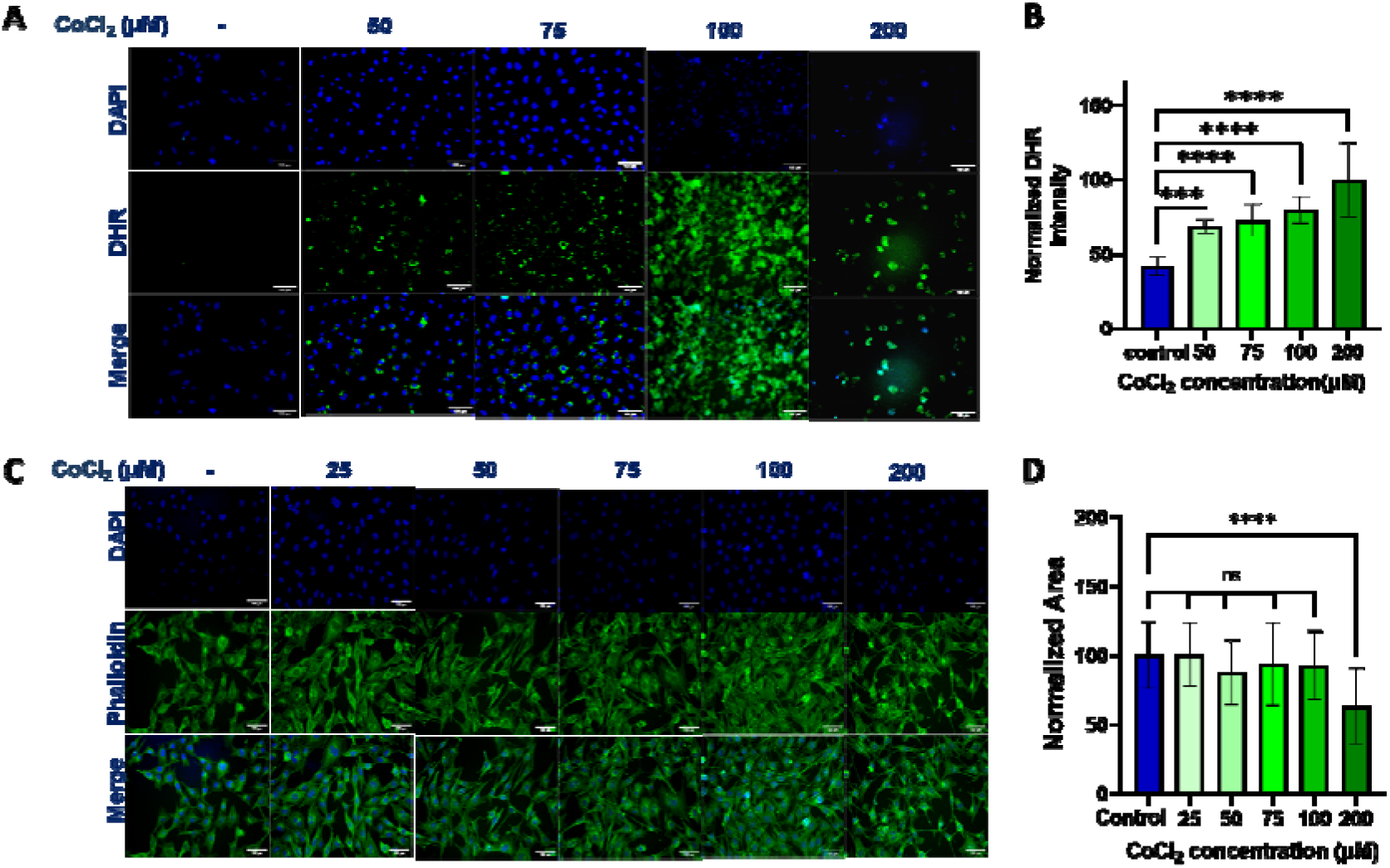
Optimization of CoCl_2_ concentration for induction of hypoxia. MDA-MB-231 cells were seeded, and hypoxia was induced with different concentrations of CoCl_2_. (A) The live cells were treated with DHR (green, to measure the ROS generation) and fixed and stained with DAPI (blue, to stain the nucleus), 63X magnification. Scale bar represents 100μm. (B) Quantification of fluorescence intensity of DHR due to the generation of ROS species. Histobars represent the mean **±** S.D. of normalized fluorescence intensity of 30 cells. (C) Confocal images of cells stained with phalloidin (green, to stain F-actin) and DAPI before and after CoCl_2_ exposure. (D) Quantification of cell area and it represents the normalized cell area. Histobars represent the mean **±** S.D. of normalized cell area of 30 cells. Asterisks denote significant difference from the control, with * p ≤ 0.05, ** p ≤ 0.01, *** p ≤ 0.001, **** p ≤ 0.0001 and ns represents non-significant difference from control.

### Optimization of hypoxic condition for MDA-MB-231 cell lines

To study the cellular effects of hypoxia, hypoxia must be induced at maximum level with minimal cell death. Therefore, there was a need to optimize the conditions. The cells were treated with different concentrations of CoCl_2_. With DHR staining, the ROS generation was monitored, and with phalloidin that stains F-actin, the cell morphology in terms of the area was observed. **Figure 1C** represents the dose-dependent effect of CoCl_2_ in the cell area, and **Figure 1A** represents the dose-dependent effect of CoCl_2_ in the ROS generation. As the CoCl_2_ concentration increases, the fluorescent intensity of DHR increases (**Figure 1B)**, indicating increased ROS generation. The cell area significantly decreased at 200μM of CoCl_2_ (**Figure 1D)**. To further confirm the cellular viability at different CoCl_2_ concentrations MTT assay was performed (**Figure S1**). With the series of CoCl_2_ concentrations, cellular death was observed more in 200μM and 400μM. Hence the optimal concentration of CoCl_2_ to induce hypoxia in MDA-MB-231 cell lines was found to be 100μM. For further studies, 100μM was used as the working concentration to induce hypoxia.

### Effect of hypoxia in cellular endocytic pathways

To study the effects of hypoxia on the dynamics of endocyticpathways in cancer cells, different endocytic pathways were tracked using fluorescently labeled cargoes such as transferrin-A488, galectin-A647, and FITC-dextran to mark CME, CIE, and FPE, respectively.**Figure 2A** shows the uptake of transferrin, galectin, and dextran in normoxic and hypoxic conditions after 24 h of hypoxia induction. After 24 h of induction of hypoxia in MDA-MB-231 cells, there was no significant change in the transferrin uptake, but there was a significant decrease in galectin uptake. Fluid phase endocytosis showed a slight significant increase in hypoxic conditions (**Figure 2B, C, D**). We initially hypothesized that there would be increased uptake via CME, CIE, and fluid-phase endocytosis in hypoxic condition to elevate nutrient uptake, thereby strengthening cellular resilience and enhancing cellular invasion. However, there was decreased uptake through CIE, no significant change inthe uptake of transferrin through CME,and a minor significant increase in FPE of dextran. The decreased uptake might benefit the cell in terms of energy preservation after 24 h of hypoxia induction. Hence, we decided to further study the change in the pattern of endocytosis after 30 mins of induction of hypoxia. **Figure 2E** shows the uptake of transferrin, galectin, and dextran in normoxic and hypoxic conditions after 30 mins of hypoxia induction. Here we observed a significant increase in uptake of galectin-3 and FITC-dextran through CIE and FPE, respectively (**Figure 2 G, H**). There was no significant change in the uptake of transferrin (**Figure 2F**). Upon short duration of induction of hypoxia, there was increased uptake via CIE and FPE; and at the later hours of hypoxia, there was a significant decrease of the same. The initial enhanced uptake of endocytosis would be because of the restoration of membrane tension when the cells were exposed to hypoxic. Also, another possible reason would be to strengthen cellular resilience by elevating nutrient uptake. Overtime decrease in endocytic uptake would be to conserve energy in the hypoxia exposed cells.

**Figure 2.**
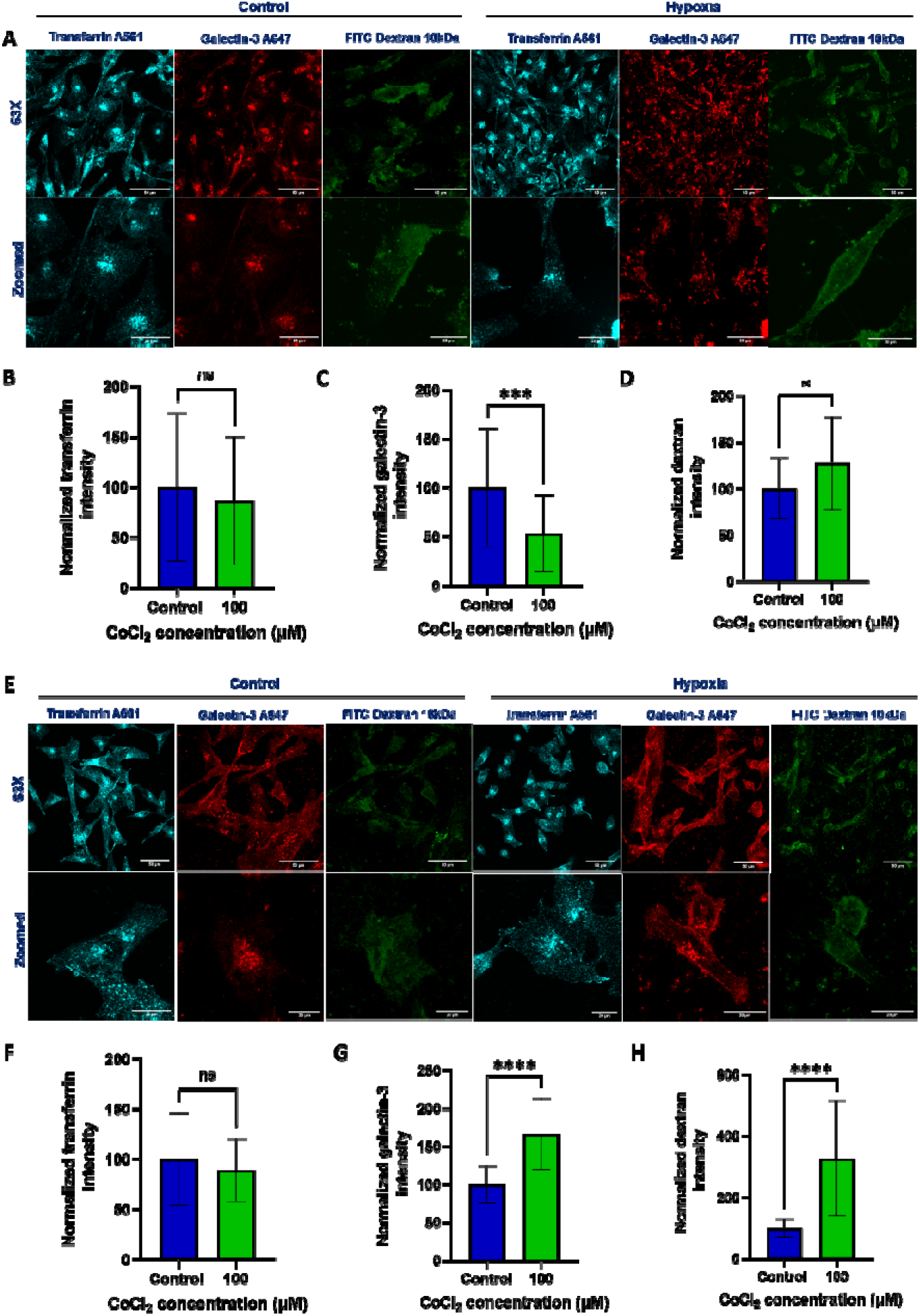
Effect of hypoxia induced by CoCl_2_ on endocytosis in MDA-MB-231 cell line: MDA-MB-231 cells were seeded, and hypoxia was induced with 100μM of CoCl_2_. (A) After 24 h of CoCl_2_ treatment, the live cells were treated with transferrin A561 (Cyan, to mark CME pathway), galectin-3 A647 (red, to mark CIE pathway), and FITC dextran 10kDa (green, to mark FPE pathway) for 15 mins. Then the cells were fixed and visualized with 63X magnification. Scale bar represents 50 μm and 20μm for the zoomed images. (B) Quantification of fluorescence intensity of the transferrin uptake after 24 h of CoCl_2_ treatment (C) Quantification of fluorescence intensity of galectin-3 uptake after 24 h of CoCl_2_ treatment. (D) Quantification of fluorescence intensity of dextran uptake after 24 h of CoCl_2_ treatment. Histobars represent the mean **±** S.D. of normalized fluorescence intensity of 30 cells. (E) After 30 mins of CoCl_2_ treatment, the live cells were treated with transferrin A561 (Cyan), galectin-3 A647 (red), and FITC dextran 10kDa (green) for 15 mins. Then the cells were fixed and visualized with 63X magnification. Scale bar represents 50 μm and 20μm for the zoomed images. (F) Quantification of fluorescence intensity of the transferrin uptake after 30 mins of CoCl_2_ treatment (G) Quantification of fluorescence intensity of galectin-3 uptake after 30 mins of CoCl_2_ treatment. (H) Quantification of fluorescence intensity of dextran uptake after 30 mins of CoCl_2_ treatment. Histobars represent the mean **±** S.D. of normalized fluorescence intensity of 30 cells. Asterisks denote significant difference from the control, with * p ≤ 0.05, ** p ≤ 0.01, *** p ≤ 0.001, **** p ≤ 0.0001 and ns represents non-significant difference from control.

### Effect of hypoxia on the sub-cellular organelles

Most pathways and organelles within cells are interconnected, and hypoxia triggers the subsequent changes in other organelles and cellular processes. Mitochondria and endoplasmic reticulum (ER) are the two essential organelles that respond dramatically to cellular stress, such as hypoxia. We checked the changes in morphology and intensity of mitochondria and ER upon hypoxia induction. **Figure 3A** represents the effect of hypoxia in the mitochondria, and **Figure 3C** represents the effect of hypoxia in the ER. Upon exposure to hypoxia for 30 mins, the cellular mitochondria intensity drastically increased when compared to the normoxic condition indicating the change in membrane environment of mitochondria upon hypoxia induction leading to secretion of more ROS (**Figure 3B**). Mitochondria appeared small, fragmented, and more intense upon hypoxia induction than controls (**Figure 3A**).

**Figure 3.**
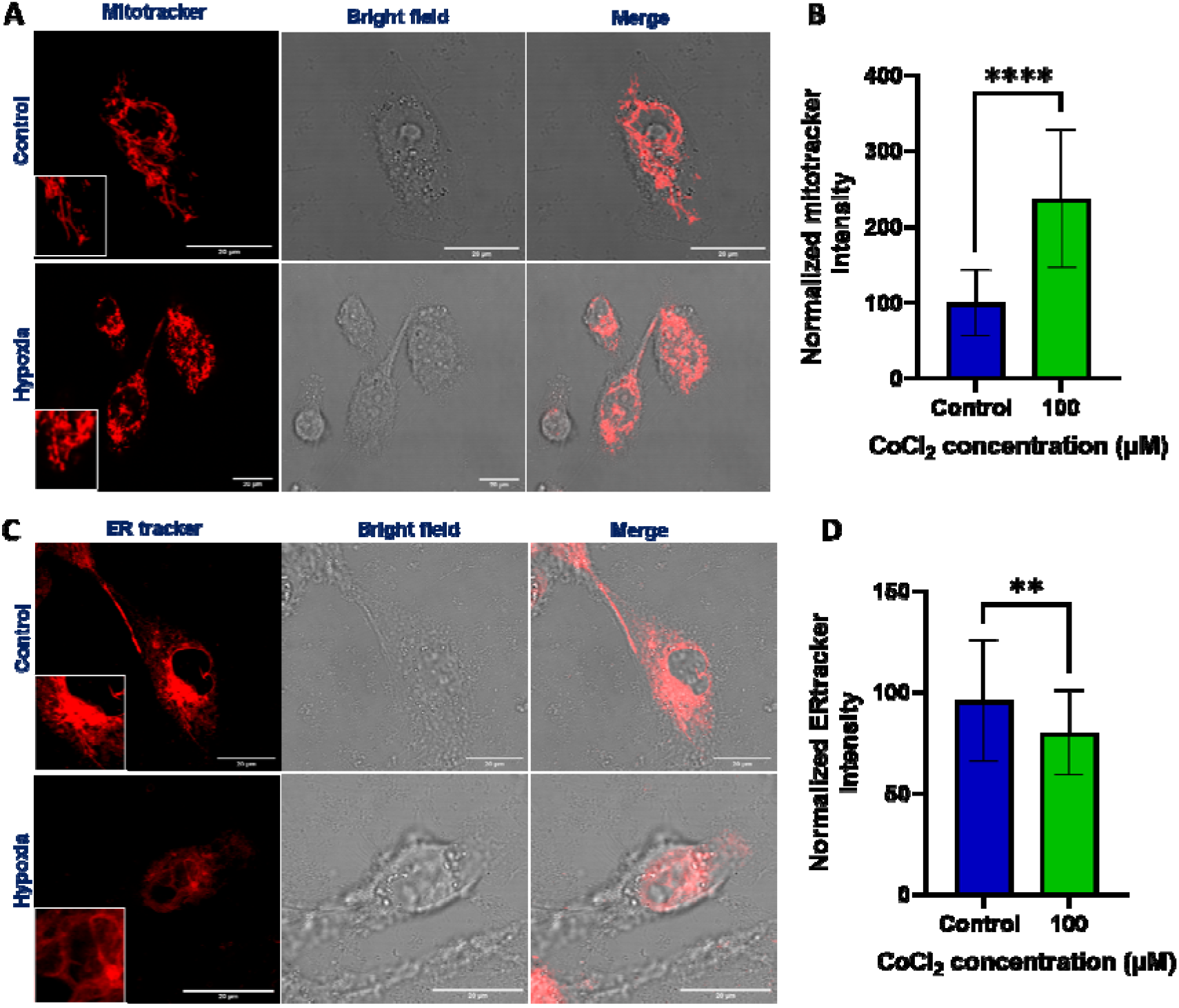
Effect of hypoxia induced by CoCl_2_ on mitochondria and endoplasmic reticulum in MDA-MB-231 cell line after 30 mins of hypoxia induction. MDA-MB-231 cells were seeded, and hypoxia was induced with 100μM of CoCl_2_. (A) After 30 mins of CoCl_2_ treatment, the live cells were treated with Mitotracker deep red FM for 30 mins. Then the cells were washed and imaged in live condition with 63X objective. Scale bar represents 20μm. (B) Quantification of fluorescence intensity of mitotracker. Histobars represent the mean **±** S.D. of normalized fluorescence intensity of 30 cells. (C) After 30 mins of CoCl_2_ treatment, the live cells were treated with ERtracker for 30 mins. Then the cells were washed and imaged in live condition with 63X objective. Scale bar represents 20μm. B) Quantification of fluorescence intensity of ER tracker. Histobars represent the mean **±** S.D. of normalized fluorescence intensity of 30 cells. Asterisks denote significant difference from the control, with * p ≤ 0.05, ** p ≤ 0.01, *** p ≤ 0.001, **** p ≤ 0.0001 and ns represents non-significant difference from control

Similarly, ER showed a decrease in intensity upon hypoxia induction (**Figure 3D**),and the network-like structures seen in the normoxic conditions were not observed in the hypoxiainduced cells (**Figure 3C**). We also tested other organelles like Golgi, peroxisomes, and lysosomes distribution and intensities upon hypoxia induction. However, those organelles did not exhibit any significant change in their morphology or intensity upon hypoxia treatment.

### Assessment of motility in normoxic and hypoxic conditions with scratch assay

Scratch assay was used to estimate the effect of hypoxia on the collective cell migration of MDA-MB-231 breast cancer cell lines(25). **Figure 4A** shows the scratch-made over a confluent monolayer monitored over 24 h in normal (left) and hypoxic (right) conditions. The gap was closed entirely at the 24^th^ hour in hypoxic condition. The images were processed, and the width and area of the wound were determined. We found that the percentage of wound closure and the rate of cell migration was significantly higher in hypoxic condition when compared with normoxic control (**Figure 4B, 4C**). Percentage wound closure was 95.05% for hypoxic condition and 68.9% for normoxic condition (**Figure 4C**). The migration rate of the cells in hypoxic and normoxic conditions was determined to be 11.37μm/h and 8.08μm/h, respectively (**Figure 4B**).

**Figure 4.**
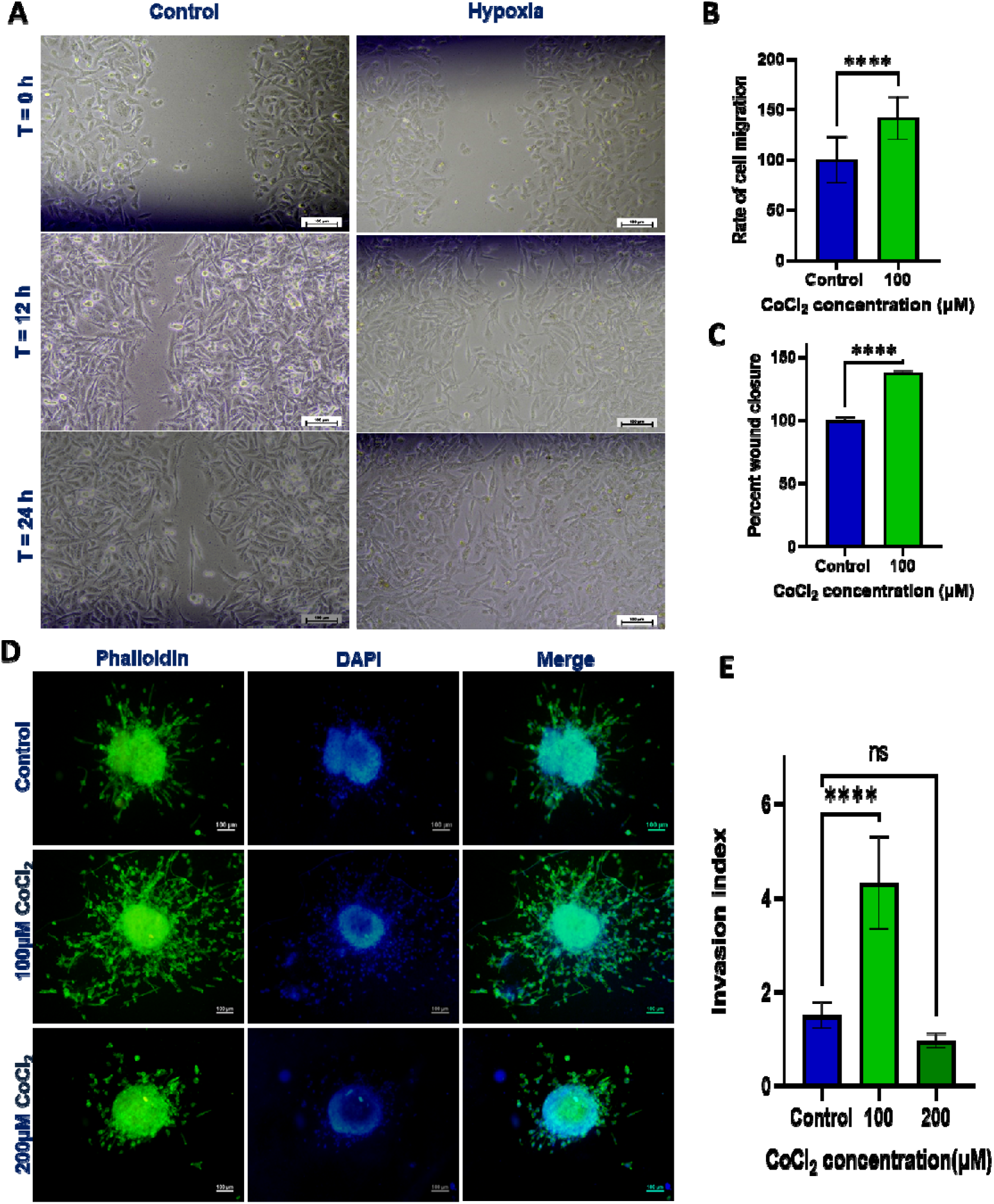
Effect of hypoxia induced by CoCl_2_ on cell migration, invasion, and proliferation in MDA-MB-231 cell line. (A) A scratch was made through a confluent monolayer of MDA-MB-231 cells using a 10μl pipette tip, and the width and area were measured 0^th^ h and 24^th^ h of CoCl_2_ treatment at 100μM concentration. Phase-contrast images represent the MDA-MB-231 monolayer scratch at 0^th^, 12^th^, and 24^th^ h in normal (left) and hypoxic (right) conditions. Scale bar represents 100μm. (B) Quantification of cell migration by measurement of the wound width.Histobars represent the mean **±** S.D. of the rate of cell migration of 60 determination of 4 independent samples. (C) Quantification of percent wound closure by measurement of the wound area. Histobars represent the mean **±** S.D. of percent wound closure of 4 independent determinations. (D) MDA-MB-231 spheroids were formed by the hanging drop method, and hypoxia was induced with CoCl_2_ concentrations of 100 and 200μM. The spheroids were fixed and stained with phalloidin (green, to stain F-actin cytoskeleton) and DAPI (blue, to stain the nucleus), 10X magnification. Scale bar represents 100μm. (E) Quantification of the spheroid invasion and represented as invasion index. Histobars represent the mean **±** S.D. of the invasion index of 5 spheroids. Asterisks denote significant difference from the control, with * p ≤ 0.05, ** p ≤ 0.01, *** p ≤ 0.001, **** p ≤ 0.0001 and ns represents non-significant difference from control.

### Effect of hypoxia on cellular invasion in the 3D spheroid model

MBA-MB-231 spheroids were formed by the hanging drop method. The spheroids were transferred to the collagen matrix to check the cellular invasion. Hypoxia was induced with CoCl_2_, and after 24 h, the spheroids were stained with phalloidin and DAPI. **Figure 4D** shows the invasion of MDA-MB-231 spheroids at normoxic and hypoxic conditions. Since MDA-MB-231 triple-negative cancer cells are inherently invasive(30), the control group of spheroids showed a considerable amount of invasion with an invasion index of 1.5 **±** 0.25. When hypoxia was induced with 100μM of CoCl_2_, the cellular invasion of spheroids was significantly higher when compared with the control group with an invasion index of 4.31 ± 0.88. However, when hypoxia was induced with 200μM of CoCl_2_, there was no significant increase in the cellular invasion of spheroids when compared with the control group (invasion index: 0.95 **±** 0.13) (**Figure 4E**). It can be attributed to cellular toxicity of high CoCl_2_ concentration. Thus, it can be concluded that hypoxia enhances cellular invasion.

### Effect of hypoxia on cell proliferation

The effect of hypoxia on cell proliferation was assessed using the MTT assay. **Figure S2** shows the effect of hypoxia in cellular proliferation. It was observed that there was a decrease in absorbance in hypoxic condition, which corresponds to decreased live cell population, which indicates decreased proliferation in the cancerous cell that experiences hypoxic condition compared to the normal cancerous cell. Cell proliferation was assessed using manual counting with a hemocytometer to validate these results. The cell count significantly decreased in hypoxia compared to normoxic cancer cells. These results revealed that the cell proliferation decreased in hypoxic condition, and it complemented the results MTT assay (Results provided in supplementary **Figure S3**).

## Conclusions

Hypoxia is a condition due to poor and improper neo-vascularisation, and the cells in those regions are deprived of nutrients; therefore, further questions arise – whether cancer cells in the hypoxic region invade and migrate more than the normal cancer cells? Whether there are changes in the cellular organelles? Whether the endocytosis pattern changes? The present study attempted to explore these questions. The CIE decreased on prolonged exposure to hypoxia, but with the shorter hypoxic duration, the CIE significantly increased.

Along with CIE, FPE increased significantly in hypoxic conditions. However, there were no significant changes observed in CME. The mitochondria in the cell drastically increased on exposure to hypoxia.We found that hypoxic cancer cells have higher migration than normal cancer cells from the scratch assay. Also, MDA-MB-231 spheroids in the 3D model have a higher invasion in the hypoxic condition than the normoxic condition. All these data suggest that when the cell endure the hypoxic stress, at the initial stage of hypoxia, there is an increase in mitochondrial membrane changes, which could be attributed to increased ROS production and increased endocytosis that could promote invasion. However, as the hypoxia prolongs, the cells may enter into the mode to conserve energy. As a result, there is decreased endocytosis and proliferation but increased migration and invasion. Though the results presented here are preliminary and no mechanistic details are present, our initial findings suggest that cellular stress like hypoxia modulates the cellular endocytic process coupled to change in mitochondrial and ER patterns that might result in decreased proliferation increased invasion and migration rates.

## Acknowledgments

We sincerely thank all the members of the DB group for critically reading the manuscript and their valuable feedback. HNA thank IITGN-MHRD, GoI MTech fellowship,and IITGN for additional fellowship. HNA specially thank Guduru Aditya Teja for valuable assistance in data acquisition. D.B. thanks SERB, GoI for Ramanujan Fellowship, IITGN, for the startup grant, DBT-EMR, Gujcost-DST, GSBTM, BRNS-BARC, and HEFA-GoI for research grants.Imaging facilities of CIF at IIT Gandhinagar are acknowledged. The authors declare no conflict of interest.

## Supplementary data

**Figure S1.**
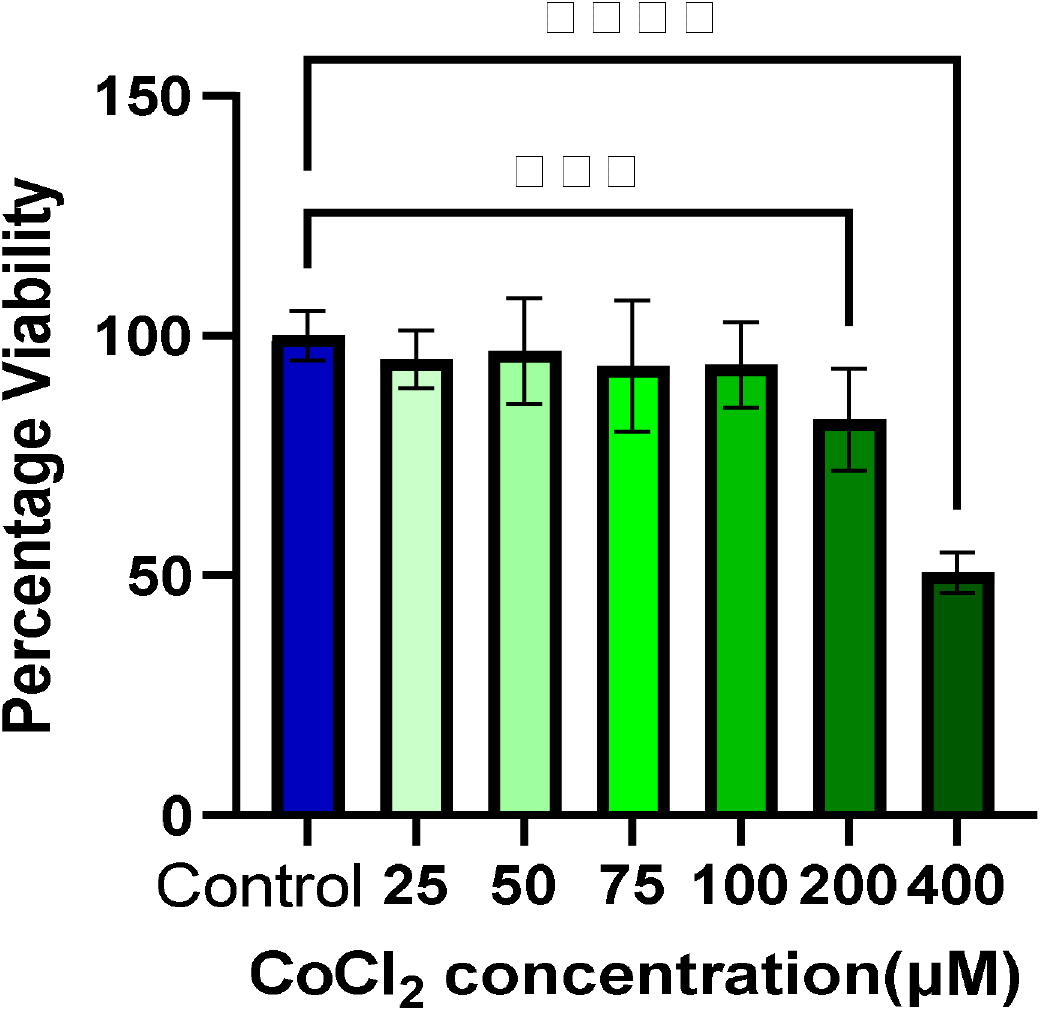
Assessment of CoCl_2_ concentration for toxicity in MDA-MB-231 cells (Optimization of CoCl_2_ concentration): MDA-MB-231 cells were seeded, and hypoxia was induced with different concentrations of CoCl_2_ The live cells were treated with MTT, and the formazan crystals formed were dissolved with DMSO. Absorbance was measured at 570nm. Histobars represent the mean ± S.D. of percentage viability of 16 independent determinations (for 25, 50, 75, 100, and 200μM) and 8 independent determination (for control and 400μM). Asterisks denote significant difference from the control, with * p ≤ 0.05, ** p ≤ 0.01, *** p ≤ 0.001, **** p ≤ 0.0001 and ns represents non-significant difference from control.

**Figure S2.**
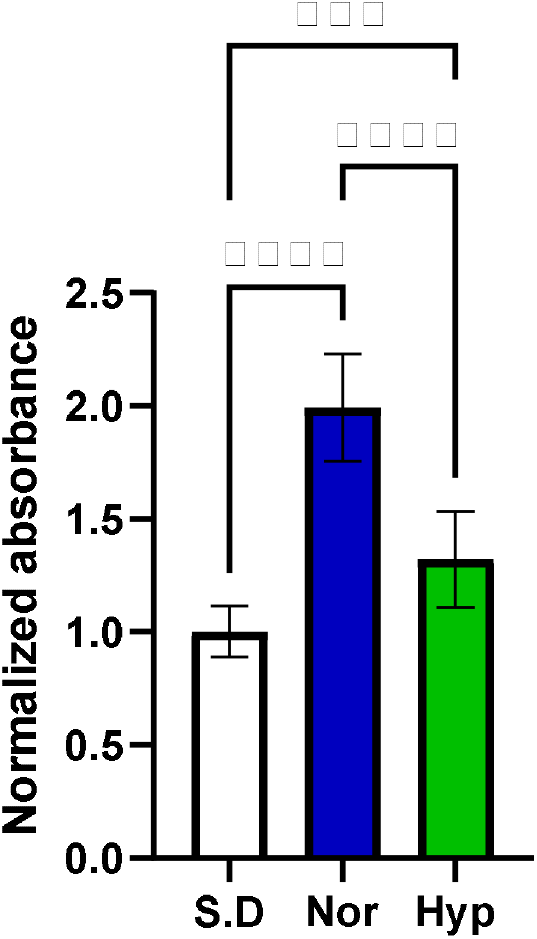
Effect of hypoxia induced by CoCl_2_ on proliferation in MDA-MB-231 cell line (MTT assay). MDA-MB-231 cells were seeded, and after 48 h of hypoxic exposure,the MTT assay was performed to assess the cell proliferation. Absorbance was measured at 570nm. Histobars represent the mean **±** S.D. of normalized absorbance of 16 independent determinations. (S.D represents the seeding density condition, Nor represents the normoxic condition, and Hyp represents hypoxic condition). Asterisks denote significant difference from the control, with * p ≤ 0.05, ** p ≤ 0.01, *** p ≤ 0.001, **** p ≤ 0.0001 and ns represents non-significant difference from control.

**Figure S3.**
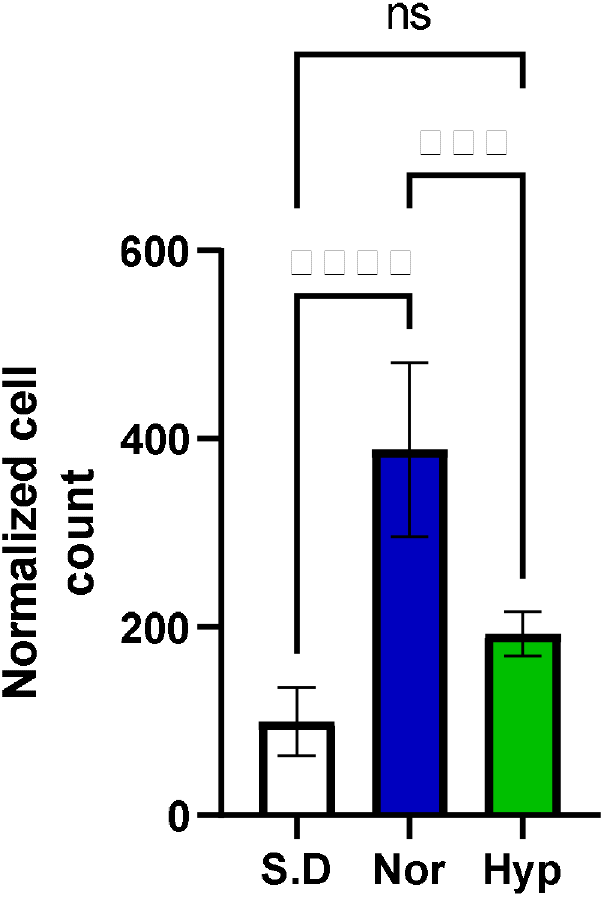
Effect of hypoxia induced by CoCl_2_ on proliferation in MDA-MB-231 cell line (Manual counting). MDA-MB-231 cells were seeded and after 48 h of hypoxic exposure the cells were trypsinized and manual cell counting was performed to assess the cell proliferation and validate the MTT result. Histobars represents the mean **±** SD of normalized cell count of 4 independent determinations. (S.D represents the seeding density condition, Nor represents normoxic condition and Hyp represents hypoxic condition). Asterisks denote significant difference from the control, with * p ≤ 0.05, ** p ≤ 0.01, *** p ≤ 0.001, **** p ≤ 0.0001 and ns represents non-significant difference from control.

